# The Impact of Vitamin A and Carotenoids on the Risk of Tuberculosis Progression

**DOI:** 10.1101/095232

**Authors:** Omowunmi Aibana, Molly F. Franke, Chuan-Chin Huang, Jerome T. Galea, Roger Calderon, Zibiao Zhang, Mercedes C. Becerra, Emily R. Smith, Alayne G. Ronnenberg, Carmen Contreras, Rosa Yataco, Leonid Lecca, Megan B. Murray

**Affiliations:** Division of General Internal Medicine, The University of Texas Health Science Center at Houston, McGovern Medical School, Houston, Texas, USA; Division of Infectious Diseases, The Miriam Hospital, Warren Alpert School of Medicine at Brown University, Providence, Rhode Island, USA; Department of Global Health and Social Medicine, Harvard Medical School, Boston, Massachusetts, USA; Department of Anesthesiology, Perioperative and Pain Medicine, Brigham and Women’s Hospital, Harvard Medical School, Boston, Massachusetts, USA; Partners In Health, Socios En Salud Sucursal Peru, Lima, Peru; Division of Global Health Equity, Brigham and Women’s Hospital, Harvard Medical School, Boston, Massachusetts, USA; Department of Global Health and Population, Harvard T. H. Chan School of Public Health, Boston, Massachusetts, USA; Department of Nutrition, University of Massachusetts Amherst, Massachusetts, USA

## Abstract

**Background:** Low and deficient levels of vitamin A are common in low and middle income countries where tuberculosis burden is high. We assessed the impact of baseline levels of vitamins A and carotenoids on TB disease risk.

**Methods and Findings:** We conducted a case-control study nested within a longitudinal cohort of household contacts of pulmonary TB cases in Lima, Peru. We screened all contacts for TB disease at 2, 6, and 12 months after enrollment. We defined cases as HIV-negative household contacts with blood samples who developed TB disease at least 15 days after enrollment of the index patient. For each case, we randomly selected 4 controls from among contacts who did not develop TB disease, matching on gender and year of age. We used conditional logistic regression to estimate odds ratios (ORs) for incident TB disease by vitamin A and carotenoids levels, controlling for other nutritional and socioeconomic factors.

Among 6751 HIV-negative household contacts with baseline blood samples, 192 developed secondary TB disease during follow-up. We analyzed 180 cases with viable samples and 709 matched controls. After controlling for possible confounders, we found that baseline vitamin A deficiency was associated with a 10-fold increase in risk of TB disease among household contacts (aOR 10.42; 95% CI 4.01–27.05; p < 0.001). This association was dose-dependent with stepwise increases in TB disease risk with each decreasing quartile of vitamin A level. Carotenoid levels were also inversely associated with TB risk among adolescents.

Our study is limited by the one year duration of follow up and by the relatively few blood samples available from household contacts under ten years of age.

**Conclusions:** Vitamin A deficiency strongly predicted risk of incident TB disease among household contacts of TB patients. Vitamin A supplementation among individuals at high risk of TB may provide an effective means of preventing TB disease.

## Introduction

Although tuberculosis (TB) incidence and mortality have fallen steadily over the past 150 years in Europe and North America, TB remains one of the top global causes of mortality from an infectious disease. The World Health Organization (WHO) reported 10.4 million new cases and 1.8 million TB deaths in 2015, the vast majority of which occurred in low- and middle-income countries [1]. In high-income countries, TB declined prior to the availability of either Bacillus Calmette–Guérin (BCG) vaccine or effective chemotherapy, and observers attribute this trend to improvements in socioeconomic status (SES) [2]. Country-level data support this hypothesis; Janssens and Rieder found an inverse linear association between per capita GDP and TB incidence [3], and Dye et al. noted that a nation’s human development index strongly predicted changes in TB incidence over time [4].

Multiple lines of evidence suggest that the association between SES and TB may be mediated through nutritional status. Studies in both humans and animals demonstrate a link between undernutrition and TB [5]. *In vitro* studies also provide insight into specific anti-mycobacterial mechanisms mediated by micronutrients [6,7]. Although multiple studies document micronutrient deficiencies in people with TB disease [8–10], few prospective studies have assessed preexisting nutritional status as a determinant of progression from TB infection to disease [11–15]. Notably, studies of vitamin A and TB have generally been conducted in populations at low risk for both nutritional deficits and TB and have been inconclusive [14,15].

Despite the lack of data on TB risk, multiple previous studies support the role of vitamin A in both innate and acquired immune responses to infection. Based on a series of studies they conducted in both animals and humans, Mellanby and Green first proposed that vitamin A is an “anti-infective” agent in the 1920s [16–18]. Subsequently, in one of the earliest reported clinical trials, Ellison showed that vitamin A supplementation reduced mortality in children with measles by 58% [19]. Current evidence demonstrates that vitamin A deficiency contributes to mortality from diarrheal diseases and measles, accounting for 1.7% of all deaths in children younger than five in low and middle income countries [20].

Here, we assessed the association between baseline levels of vitamin A and carotenoids and the risk of TB disease among household contacts of patients with pulmonary TB in Lima, Peru.

## Methods

The study was approved by the Institutional Review Board of Harvard School of Public Health and the Research Ethics Committee of the National Institute of Health of Peru. All participants or guardians provided written informed consent.

### Setting and Study Design

We conducted a case-control study nested within a prospective longitudinal cohort of household contacts (HHCs) of TB cases in Lima, Peru. The study area includes 20 districts of metropolitan Lima with a population of approximately 3.3 million residents living in urban areas and peri-urban, informal shantytown settlements. Between September 2009 and August 2012, we identified index patients 15 years and older diagnosed with pulmonary TB at 106 participating health centers. Within two weeks of enrolling an index patient, we enrolled his or her HHCs and screened for TB disease. HHCs with symptoms were referred for sputum smear microscopy and mycobacteriology culture and treatment according to Peru’s national guidelines [21]. HHCs aged 19 and under, and individuals with specified co-morbidities were offered isoniazid preventive therapy (IPT).

Upon enrollment of HHCs, we collected the following: age, gender, height, weight, BCG vaccination scars, IPT use, alcohol and tobacco use, self-reported diabetes mellitus (DM), comorbid diseases (heart disease, high blood pressure, asthma, kidney disease, use of steroids or chemotherapy or immunosuppressant, any other self-reported chronic illness), TB disease history, and housing asset information (housing type, number of rooms, water supply, sanitation facilities, lighting, composition of exterior walls and floor and roof materials). In HHCs with no prior history of TB infection or disease, we obtained a tuberculin skin test (TST). We offered HIV testing to HHCs and invited all HHCs to provide a venous blood sample; 60% of those ≥10 years old provided this sample.

All HHCs were evaluated for signs and symptoms of pulmonary and extra-pulmonary TB (EPTB) disease at two, six, and 12 months after enrollment. We considered HHCs to have developed incident secondary TB disease if the diagnosis was confirmed by a health center physician more than 14 days after index case enrollment and with co-prevalent disease if they were diagnosed earlier. We defined secondary TB disease in HHCs under age 18 according to consensus guidelines for diagnosis of TB disease in children [22].

For this study, we defined “cases” to be HIV-negative HHCs who developed incident secondary TB disease during one year of follow-up. We randomly selected four controls for each case from among HHCs who did not develop TB disease within one year, matching on gender and age by year.

### Laboratory Methods

We stored serum samples at −80°C after collection until analysis at end of study follow-up. Laboratory personnel were unaware of the specimens’ case-control status. All samples were handled identically and assayed randomly. Levels of retinol and carotenoids (lutein+zeaxanthin,β-cryptoxanthin, total lycopene, α-carotene, and total β-carotene) were measured by high-performance liquid chromatography [23]. The interassay coefficients of variation for retinol and carotenoids were < 5%. We also measured levels of total 25–(OH)D with a commercial competitive enzyme immunoassay kit (Immunodiagnostic Systems Inc., Fountain Hills, AZ). The coefficient of variation at different levels of 25–(OH)D ranged from 4.6% to 8.7%.

### Statistical Analysis

We defined vitamin A deficiency (VAD) as serum retinol < 200 μg/L (0.70 μmol/L) [24] and vitamin D deficiency as serum 25–(OH)D < 50 nmol/L, insufficiency as 50–75 nmol/L and sufficiency as > 75nmol/L [11,25]. We calculated body mass index (BMI) as weight in kilograms divided by square of height in meters. For children and adolescent HHCs < 20 years, we used WHO age and gender-specific BMI z-scores tables to classify those with BMI z-score < −2 as underweight and those with z-score >2 as overweight [26]. For adults ≥ 20 years, we classified nutritional status as: underweight (BMI < 18.5 kg/m^2^), normal (BMI 18.5–< 25 kg/m^2^), and overweight (BMI ≥ 25 kg/m^2^). We classified HHCs as heavy drinkers if they drank ≥ 40g or ≥ 3 alcoholic drinks daily based on past year recall. We derived an SES score using principal components analysis of housing asset index weighted by household size [27]. We considered HHCs infected with TB at baseline if they reported previous TB disease or a positive TST or had a TST ≥ 10 mm at enrollment.

We constructed conditional logistic regression models to evaluate the association between quartiles of vitamin A and carotenoids and risk of TB disease. We performed tests for linear trend across quartiles. Multivariable-adjusted models included baseline covariates identified a priori as potential confounders (BMI, SES, alcohol and tobacco use, receipt of IPT, TB disease history, comorbid disease, self-reported DM and index patient smear status). We also assessed the impact of vitamin A deficiency on incident TB disease and examined the interaction between vitamin A and D deficiency on TB disease risk. We stratified the final adjusted models by age. Under our method of control selection, odds ratios approximate risk ratios because secondary TB disease was relatively rare in this cohort [28].

In sensitivity analyses, we stratified by baseline TB infection status and separately restricted the analysis to cases (and their matched controls) diagnosed at least 90 days after index patient enrollment. Finally, we repeated our analyses for cases (and their matched controls) with microbiologically confirmed TB.

For all tests, we considered two-sided p values < .05 as statistically significant. Data were analyzed using SAS v9.4 (SAS Institute, 2013).

## Results

Among 6751 HIV-negative HHCs with blood samples, 258 HHCs developed TB disease, of which 66 were co-prevalent and 192 were secondary. Pulmonary disease accounted for 236 (91.5%) cases and extra-pulmonary for 16 (6.2%). Of 180 secondary TB cases with viable blood samples, 147 (81.7%) were microbiologically confirmed (Fig1). Table 1 shows baseline characteristics of cases and controls. Cases had lower baseline median levels of vitamin A and carotenoids compared to controls (Table 2). Although the pro-vitamin A carotenoids, β-cryptoxanthin, α-carotene, and β-carotene contribute to retinol levels, here we found they were not correlated with measured retinol levels (R 0.04 to 0.10). Table 3 demonstrates significant univariate associations between incident TB disease and baseline quartiles of vitamin A, lutein/zeaxanthin, β-cryptoxanthin, total lycopene, and α-carotene. After multivariable adjustment for matching factors and confounders, household contacts in the lowest quartile of vitamin A had seven times the risk of TB disease compared to those in the highest quartile (OR 7.03; 95% CI 3.14–15.71; P for trend <0.001) [Table 3]. Lower levels of lutein/zeaxanthin (Q1 vs Q4: OR 2.10; 95% CI 1.17–3.77; P for trend = 0.06) and β-cryptoxanthin (Q1 vs Q4: OR 2.61; 95% CI 1.41–4.86; P for trend = 0.002) were also associated with increased risk of TB disease.

**Fig 1.**
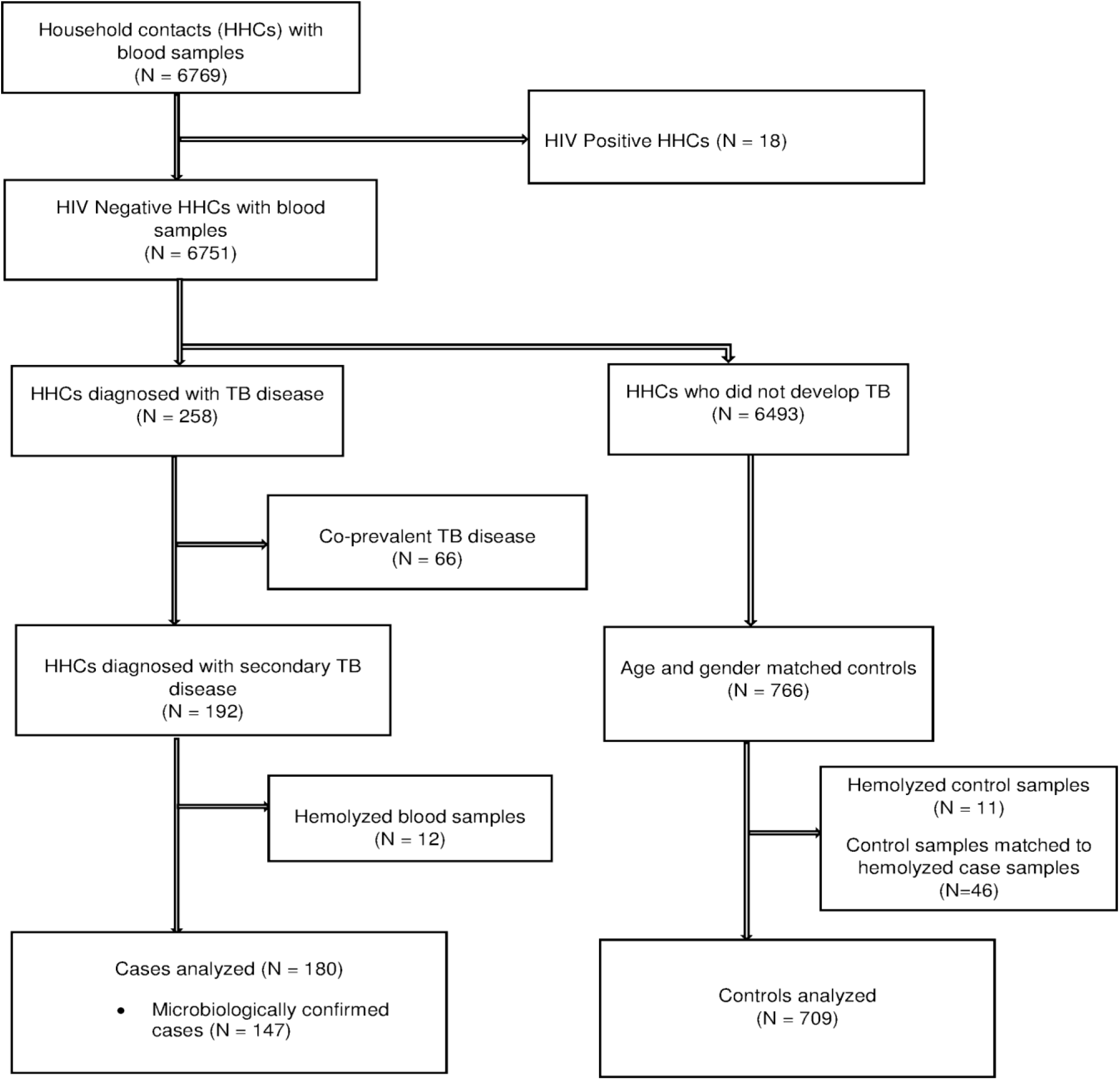
Flow Diagram for Nested Case-Control Study.

**Table 1.** Baseline Characteristics of Incident TB Cases and Matched Controls.

|  | **Cases (N = 180)**<br>**n (%)** | **N<sup>a</sup>**<br>**(Cases)** | **Control (N = 709) n**<br>**(%)** | **N<sup>a</sup>**<br>**(Controls)** |
| --- | --- | --- | --- | --- |
| Age Categories in years |  |  |  |  |
| < 10 | 4 (2.2) |  | 16 (2.3) |  |
| 10 to 19 | 50 (27.8) |  | 200 (28.2) |  |
| ≥ 20 | 126 (70.0) |  | 493 (69.5) |  |
| Male | 94 (52.2) |  | 366 (51.6) |  |
| BMI Categories |  | 179 |  | 706 |
| Underweight | 8 (4.5) |  | 6 (0.9) |  |
| Overweight | 45 (25.1) |  | 299 (42.4) |  |
| Normal | 126 (70.4) |  | 401 (56.8) |  |
| Socioeconomic Status |  | 171 |  | 698 |
| Low | 77 (45.0) |  | 228 (32.7) |  |
| Middle | 66 (38.6) |  | 326 (46.7) |  |
| High | 28 (16.4) |  | 144 (20.6) |  |
| Heavy Alcohol Use | 14 (8.1) | 174 | 64 (9.3) | 692 |
| Current Smoking | 13 (7.4) | 176 | 78 (11.2) | 699 |
| Self-Reported Diabetes | 6 (3.4) | 179 | 11 (1.6) | 701 |
| Comorbid Disease | 37 (20.6) |  | 175 (24.7) |  |
| Isoniazid Preventive Therapy | 7 (3.9) |  | 108 (15.2) |  |
| BCG scar | 159 (88.3) |  | 628 (88.6) |  |
| History of TB | 34 (18.9) |  | 55 (7.8) | 708 |
| TB Infection at baseline | 145 (82.4) | 176 | 281 (41.1) | 683 |
| **Index Patient Characteristics** |  |  |  |  |
| Smear Positive | 156 (86.7) |  | 486 (68.7) | 707 |
| Cavitary Disease | 54 (30.2) | 179 | 175 (25.1) | 697 |

BCG, Bacillus Calmette–Guérin.
<sup>a</sup>Total number of subjects with data for corresponding variable.

**Table 2.** Baseline Levels of Micronutrients Among Cases and Controls.

|  | **Cases (N = 180)**<br>**median (IQR) or n (%)** | **Control (N = 709)**<br>**median (IQR) or n (%)** |
| --- | --- | --- |
| Vitamin A (µg/L) | 316.5 (250.5 – 381.4) | 373.6 (310.4 – 460.2) |
| Vitamin A Deficient (< 200 µg/L) | 24 (13.3) | 16 (2.3) |
| 25–OH Vitamin D (nmol/L) | 53.9 (42.7 – 64.0) | 54.7 (44.5 – 67.1) |
| Vitamin D Deficient (< 50 nmol/L) | 76 (42.2) | 259 (36.5) |
| Vitamin D Insufficient (50–75 nmol/L) | 84 (46.7) | 348 (49.1) |
| Vitamin D Sufficient (> 75 nmol/L) | 20 (11.1) | 102 (14.4) |
| Lutein + Zeaxanthin (µg/L) | 150.5 (111.1 – 186.1) | 153.6 (121.5 – 201.8) |
| β-cryptoxanthin (µg/L) | 107.3 (68.2 – 196.7) | 142.3 (85.5 – 240.5) |
| Total Lycopene (µg/L) | 99.7 (64.9 – 147.3) | 112.9 (77.6 – 164.5) |
| α-carotene (µg/L) | 65.3 (44.2 – 97.1) | 75.8 (52.6 – 110.0) |
| Total-β-carotene (µg/L) | 123.2 (67.9 – 192.4) | 138.0 (86.3 – 216.4) |

**Table 3.**
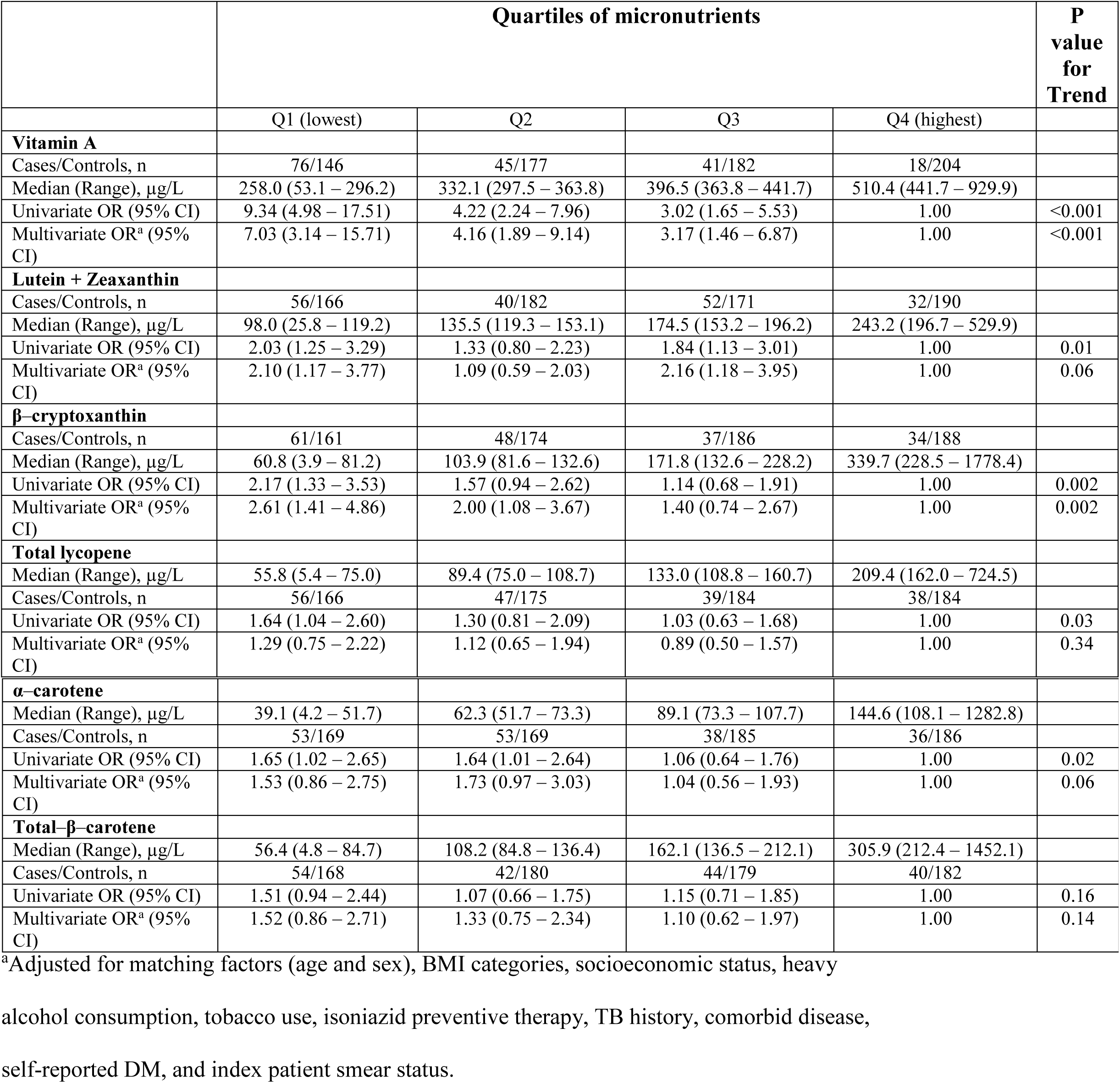
Micronutrient Levels by Quartile and the Risk of TB Disease among Household Contacts of Index TB Patients (N = 889).

|  | **Quartiles of micronutrients** |  |  |  | **P**<br>**value**<br>**for**<br>**Trend** |
| --- | --- | --- | --- | --- | --- |
|  | Q1 (lowest) | Q2 | Q3 | Q4 (highest) |  |
| **Vitamin A** |  |  |  |  |  |
| Cases/Controls, n | 76/146 | 45/177 | 41/182 | 18/204 |  |
| Median (Range), µg/L | 258.0 (53.1 – 296.2) | 332.1 (297.5 – 363.8) | 396.5 (363.8 – 441.7) | 510.4 (441.7 – 929.9) |  |
| Univariate OR (95% CI) | 9.34 (4.98 – 17.51) | 4.22 (2.24 – 7.96) | 3.02 (1.65 – 5.53) | 1.00 | <0.001 |
| Multivariate OR<sup>a</sup> (95% CI) | 7.03 (3.14 – 15.71) | 4.16 (1.89 – 9.14) | 3.17 (1.46 – 6.87) | 1.00 | <0.001 |
| **Lutein + Zeaxanthin** |  |  |  |  |  |
| Cases/Controls, n | 56/166 | 40/182 | 52/171 | 32/190 |  |
| Median (Range), µg/L | 98.0 (25.8 – 119.2) | 135.5 (119.3 – 153.1) | 174.5 (153.2 – 196.2) | 243.2 (196.7 – 529.9) |  |
| Univariate OR (95% CI) | 2.03 (1.25 – 3.29) | 1.33 (0.80 – 2.23) | 1.84 (1.13 – 3.01) | 1.00 | 0.01 |
| Multivariate OR<sup>a</sup> (95% CI) | 2.10 (1.17 – 3.77) | 1.09 (0.59 – 2.03) | 2.16 (1.18 – 3.95) | 1.00 | 0.06 |
| **β-cryptoxanthin** |  |  |  |  |  |
| Cases/Controls, n | 61/161 | 48/174 | 37/186 | 34/188 |  |
| Median (Range), µg/L | 60.8 (3.9 – 81.2) | 103.9 (81.6 – 132.6) | 171.8 (132.6 – 228.2) | 339.7 (228.5 – 1778.4) |  |
| Univariate OR (95% CI) | 2.17 (1.33 – 3.53) | 1.57 (0.94 – 2.62) | 1.14 (0.68 – 1.91) | 1.00 | 0.002 |
| Multivariate OR<sup>a</sup> (95% CI) | 2.61 (1.41 – 4.86) | 2.00 (1.08 – 3.67) | 1.40 (0.74 – 2.67) | 1.00 | 0.002 |
| **Total lycopene** |  |  |  |  |  |
| Median (Range), µg/L | 55.8 (5.4 – 75.0) | 89.4 (75.0 – 108.7) | 133.0 (108.8 – 160.7) | 209.4 (162.0 – 724.5) |  |
| Cases/Controls, n | 56/166 | 47/175 | 39/184 | 38/184 |  |
| Univariate OR (95% CI) | 1.64 (1.04 – 2.60) | 1.30 (0.81 – 2.09) | 1.03 (0.63 – 1.68) | 1.00 | 0.03 |
| Multivariate OR<sup>a</sup> (95% CI) | 1.29 (0.75 – 2.22) | 1.12 (0.65 – 1.94) | 0.89 (0.50 – 1.57) | 1.00 | 0.34 |
| **α-carotene** |  |  |  |  |  |
| Median (Range), $\mu\text{g/L}$ | 39.1 (4.2 – 51.7) | 62.3 (51.7 – 73.3) | 89.1 (73.3 – 107.7) | 144.6 (108.1 – 1282.8) | |
| Cases/Controls, n | 53/169 | 53/169 | 38/185 | 36/186 |  |
| Univariate OR (95% CI) | 1.65 (1.02 – 2.65) | 1.64 (1.01 – 2.64) | 1.06 (0.64 – 1.76) | 1.00 | 0.02 |
| Multivariate OR<sup>a</sup> (95% CI) | 1.53 (0.86 – 2.75) | 1.73 (0.97 – 3.03) | 1.04 (0.56 – 1.93) | 1.00 | 0.06 |
| **Total-β-carotene** |  |  |  |  |  |
| Median (Range), $\mu\text{g/L}$ | 56.4 (4.8 – 84.7) | 108.2 (84.8 – 136.4) | 162.1 (136.5 – 212.1) | 305.9 (212.4 – 1452.1) | |
| Cases/Controls, n | 54/168 | 42/180 | 44/179 | 40/182 |  |
| Univariate OR (95% CI) | 1.51 (0.94 – 2.44) | 1.07 (0.66 – 1.75) | 1.15 (0.71 – 1.85) | 1.00 | 0.16 |
| Multivariate OR<sup>a</sup> (95% CI) | 1.52 (0.86 – 2.71) | 1.33 (0.75 – 2.34) | 1.10 (0.62 – 1.97) | 1.00 | 0.14 |

<sup>a</sup>Adjusted for matching factors (age and sex), BMI categories, socioeconomic status, heavy alcohol consumption, tobacco use, isoniazid preventive therapy, TB history, comorbid disease, self-reported DM, and index patient smear status.

Tables 4 provides univariate and adjusted odds ratios for vitamin A deficiency. HHCs with baseline vitamin A deficiency were at 10.42-fold increased risk of incident TB disease (95% CI 4.01–27.05; P < 0.001). A statistical test for interaction between vitamins A and D deficiencies was not significant (P for interaction = 0.28).

**Table 4.**
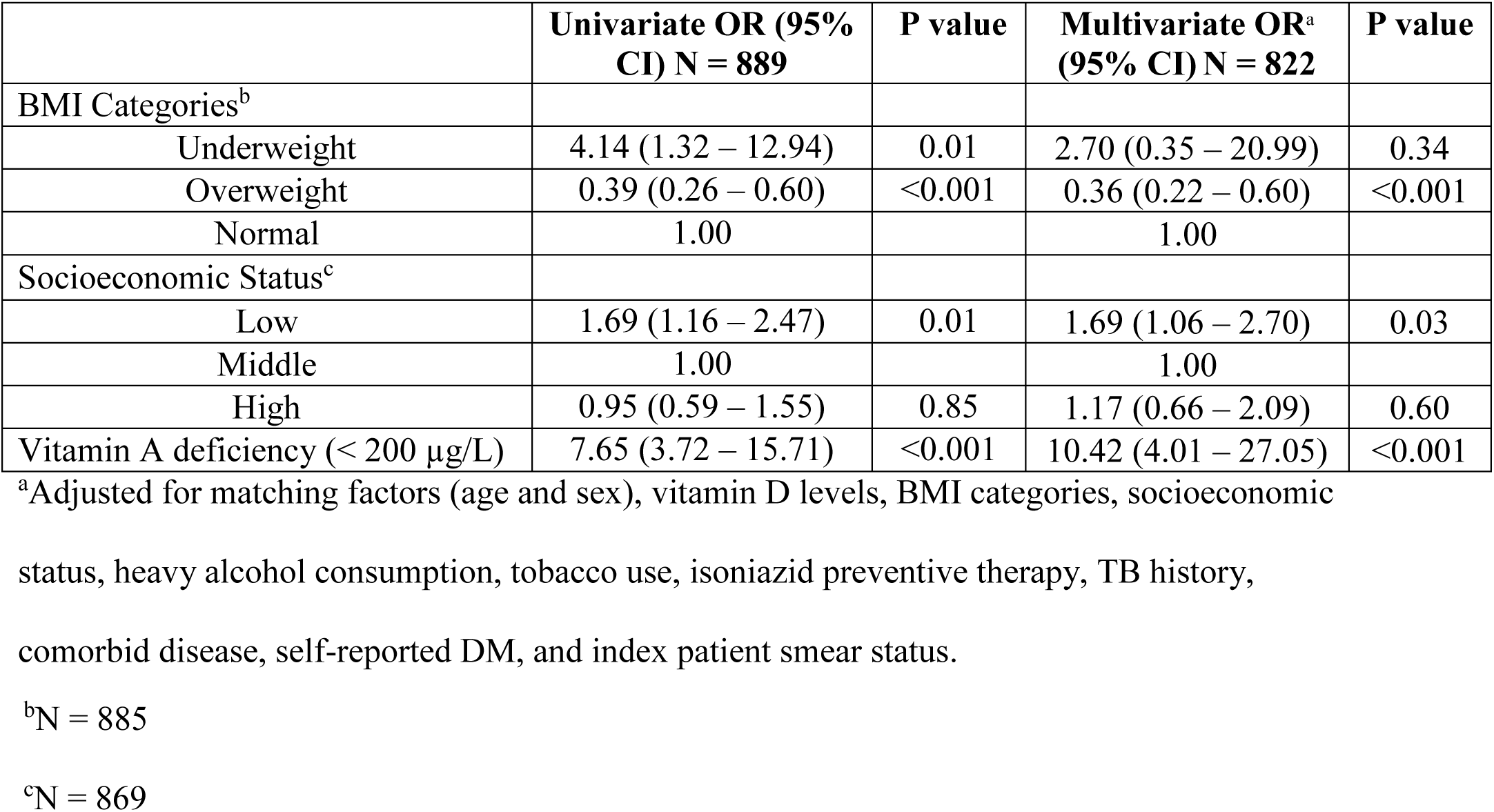
Vitamins A deficiency with Selected Risk Factors and Risk of TB Disease.

Stratification by age showed that VAD increased the risk of TB disease in 10–19 year olds nearly 20-fold (aOR 18.58; 95% CI 3.54–97.60; P = 0.001) and in ≥ 20 year olds by 10-fold (aOR 10.21; 95% CI 2.43–42.91; P = 0.002) [Table 5]. Adolescents aged 10–19 years in the lowest quartile of carotenoid levels were at higher risk for TB disease but there was no effect in older HHCs.

**Table 5.**
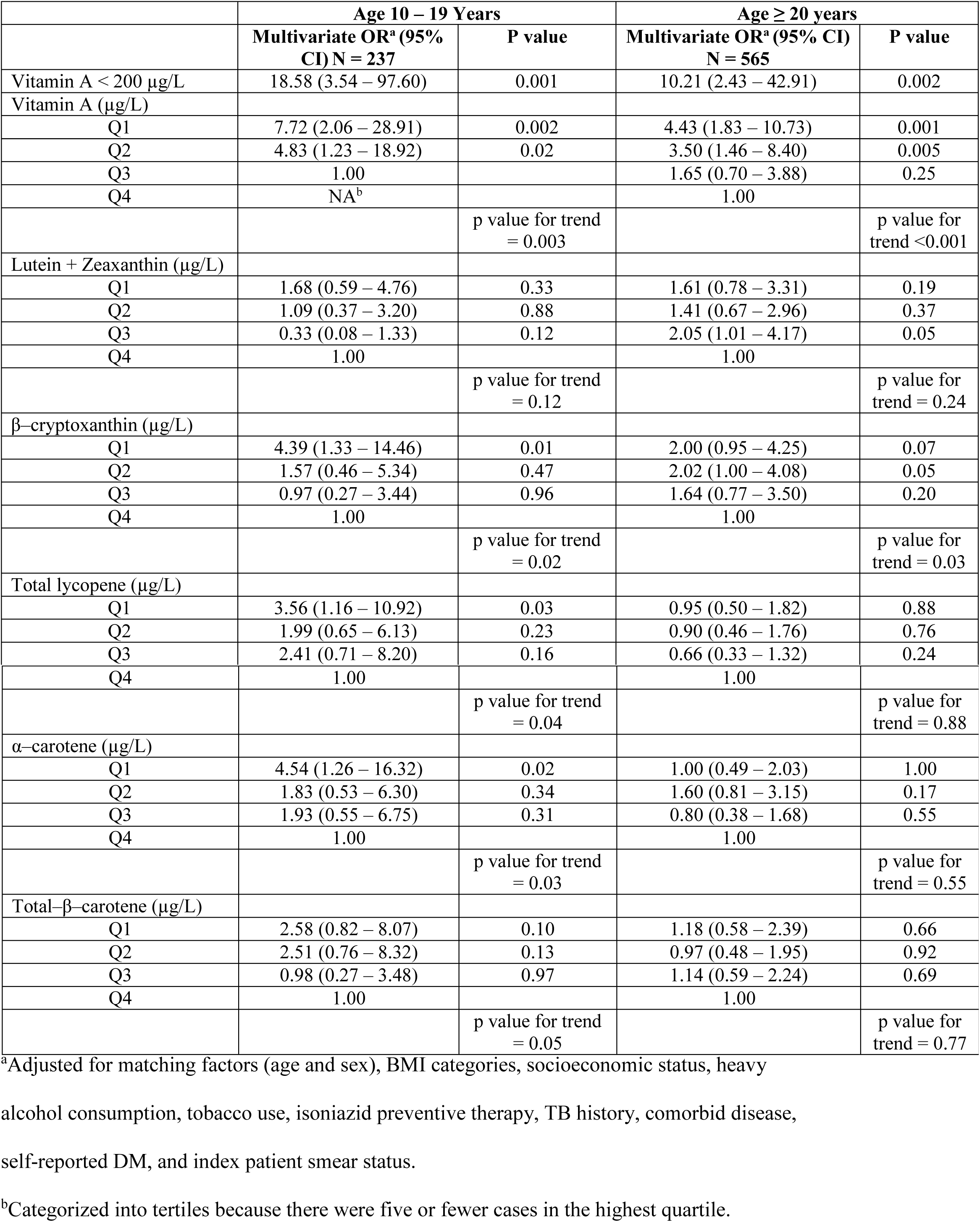
Micronutrient Levels and Risk of TB disease Stratified by Age Category.

To rule out the possibility that vitamin A deficiency in cases might have resulted from metabolic derangements due to early active TB disease that was not clinically apparent, we stratified by baseline infection status and found vitamin A levels remained strongly associated with TB disease in both groups (Table 6). In a second sensitivity analysis in which we excluded cases (and their matched controls) diagnosed within 90 days of index patient enrollment, multivariable analysis also continued to show a strong inverse association between quartiles of vitamin A and TB disease although the association with VAD was no longer statistically significant (Table 6).

**Table 6.** Vitamin A Levels and Risk of TB disease Stratified by Baseline Infection Status and for TB Diagnosed at least 90 days After Index Case Enrollment.

|  | Cases/Controls | Multivariate OR<sup>a</sup><br>(95% CI) | P value |
| --- | --- | --- | --- |
| **Negative TB Infection at baseline** |  |  |  |
| Vitamin A (µg/L) |  |  |  |
| Q1 | 12/106 | 4.38 (1.24 – 15.47) | 0.02 |
| Q2 | 13/144 | 3.71 (1.12 – 12.33) | 0.03 |
| Q3 | 6/152 | 1.00 |  |
| Q4 | NA<sup>b</sup> |  |  |
|  |  |  | p value for trend<br>= 0.02 |
| Vitamin A Deficient (<200 µg/L) | 3/8 | 20.32 (2.91 – 141.70) | 0.002 |
| **Positive TB Infection at baseline** |  |  |  |
| Vitamin A (µg/L) |  |  |  |
| Q1 | 80/90 | 2.56 (1.17 – 5.58) | 0.02 |
| Q2 | 40/89 | 1.83 (0.85 – 3.92) | 0.12 |
| Q3 | 25/102 | 1.00 |  |
| Q4 | NA |  |  |
|  |  |  | p value for trend<br>= 0.02 |
| Vitamin A Deficient (<200 µg/L) | 21/8 | 5.31 (1.63 – 17.25) | 0.01 |
| **TB diagnosed ≥ 90 days** |  |  |  |
| Vitamin A (µg/L) |  |  |  |
| Q1 | 27/70 | 4.21 (1.20 – 14.74) | 0.02 |
| Q2 | 24/73 | 5.25 (1.44 – 19.10) | 0.01 |
| Q3 | 19/79 | 2.91 (0.87 – 9.77) | 0.08 |
| Q4 | 8/89 | 1.00 |  |
|  |  |  | p value for trend<br>= 0.02 |
| Vitamin A Deficient (<200 µg/L) | 3/8 | 2.15 (0.33 – 13.80) | 0.42 |

<sup>a</sup>Adjusted for matching factors (age and sex), BMI categories, socioeconomic status, heavy alcohol consumption, tobacco use, isoniazid preventive therapy, TB history, comorbid disease, self-reported DM, and index patient smear status.
<sup>b</sup>Categorized into tertiles because there were five or fewer cases in the highest quartile.

Restricting the analysis to microbiologically confirmed cases further strengthened the association between vitamin A levels and TB disease (Table 7). We were unable to stratify by IPT use because few HHCs had received IPT.

**Table 7.**
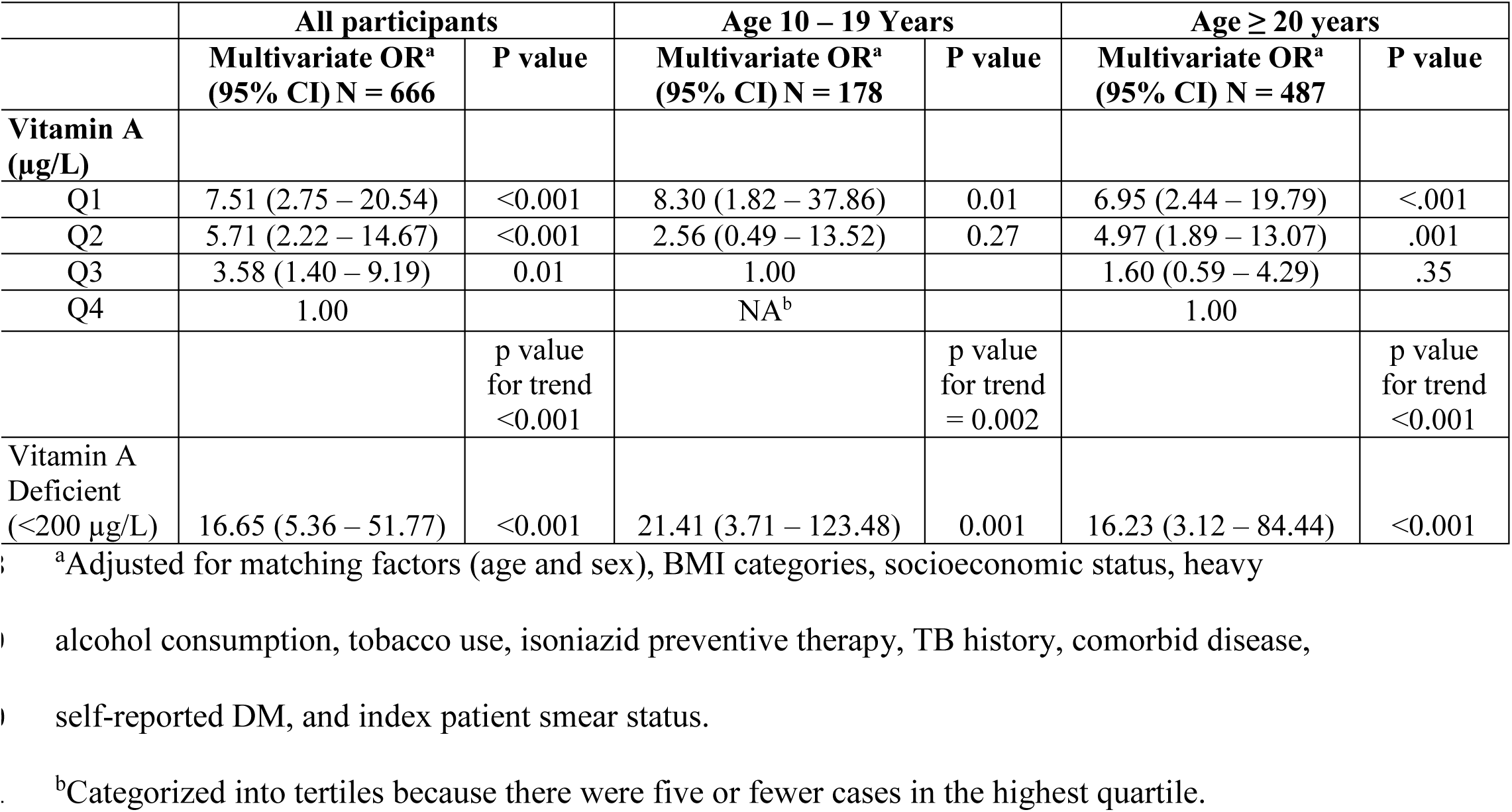
Vitamin A levels and Risk of Microbiologically Confirmed TB Disease.

|  | **All participants** |  | **Age 10 – 19 Years** |  | **Age ≥ 20 years** |  |
| --- | --- | --- | --- | --- | --- | --- |
|  | **Multivariate OR<sup>a</sup>**<br>**(95% CI) N = 666** | **P value** | **Multivariate OR<sup>a</sup>**<br>**(95% CI) N = 178** | **P value** | **Multivariate OR<sup>a</sup>**<br>**(95% CI) N = 487** | **P value** |
| **Vitamin A**<br>**(µg/L)** |  |  |  |  |  |  |
| Q1 | 7.51 (2.75 – 20.54) | <0.001 | 8.30 (1.82 – 37.86) | 0.01 | 6.95 (2.44 – 19.79) | <.001 |
| Q2 | 5.71 (2.22 – 14.67) | <0.001 | 2.56 (0.49 – 13.52) | 0.27 | 4.97 (1.89 – 13.07) | .001 |
| Q3 | 3.58 (1.40 – 9.19) | 0.01 | 1.00 |  | 1.60 (0.59 – 4.29) | .35 |
| Q4 | 1.00 |  | NA<sup>b</sup> |  | 1.00 |  |
|  |  | p value<br>for trend<br><0.001 |  | p value<br>for trend<br>= 0.002 |  | p value<br>for trend<br><0.001 |
| Vitamin A<br>Deficient<br>(<200 µg/L) | 16.65 (5.36 – 51.77) | <0.001 | 21.41 (3.71 – 123.48) | 0.001 | 16.23 (3.12 – 84.44) | <0.001 |

<sup>a</sup>Adjusted for matching factors (age and sex), BMI categories, socioeconomic status, heavy
alcohol consumption, tobacco use, isoniazid preventive therapy, TB history, comorbid disease,
self-reported DM, and index patient smear status.
<sup>b</sup>Categorized into tertiles because there were five or fewer cases in the highest quartile.

## Discussion

Here, we found that vitamin A levels strongly predict progression to TB disease in a cohort of Peruvians with household exposure to an index TB patient. This relationship remained strong after adjustment for socioeconomic status, BMI and other comorbidities that might be associated with increased TB disease risk and lower vitamin A levels. Although the association is most extreme at levels consistent with a diagnosis of VAD, we observed a dose-response relationship with significantly increased risk of TB disease even at levels usually classified as normal. Carotenoids were also inversely associated with risk of TB progression among adolescents, although the effect of individual carotenoids was less extreme than for vitamin A.

Although previous studies have documented low vitamin A levels among patients with TB disease [8,9,29,30], the causal direction of this association has been unclear. TB disease can lead to profound weight loss, and micronutrient deficiencies observed in patients often resolve with TB treatment [30]. To date, no published studies have reported on the impact of vitamin A supplementation on incident TB disease, and no clinical trials have demonstrated an impact of supplementation on TB treatment outcomes [30].

In contrast to the plethora of work on micronutrients and TB treatment outcomes [30], few studies have prospectively investigated the role of pre-existing vitamin A status in the development of TB disease. Cegielski et al. observed that less than 2% of the US-based NHANES 1 cohort [14] had baseline vitamin A levels under 300 μg/L (1.05 μmol/L), and only 61 people developed TB disease over 20 years of follow-up. Although TB incidence was nearly three-fold higher among those with low vitamin A levels, confidence intervals for the hazard ratio spanned from 0.70 to 11.4. These results are consistent with a study conducted in Philadelphia between 1942–49, in which African American men with low baseline vitamin A levels [15] were three times more likely to develop TB disease than those with higher levels. Interestingly, the mean vitamin A level was 300 μg/L (1.05 μmol/L), suggesting that the comparison was between men with low and very low levels, rather than among people with the spectrum of levels we observed in Lima. This study also found a very strong effect of vitamin C deficiency, with all those who developed TB disease having baseline vitamin C levels below 6 mg/L (34.07 μmol/L).

Multiple other lines of evidence suggest an association between vitamin A levels and TB risk. In vitro studies have demonstrated that vitamin A and its metabolites cause dose-dependent inhibition of *Mycobacterium tuberculosis* (Mtb) growth in culture [31,32]. Experimental studies in animal and cell models of TB infection also provide support for the protective role of retinol. Although early studies in various models gave conflicting results [33–35], more recent studies in mice [36], rats [37], THP-1 cells [38], and cultured human macrophages [39] show reduced Mtb bacillary loads after treatment with retinoic acid. Interestingly, the dose of retinoic acid required to produce a protective effect in human macrophages is lower when it was added prior to infection than when it was added afterwards [39].

Vitamin A also affects the immune system in various ways relevant to the pathogenesis of TB [40]. Most of its effects are exerted by retinoic acid, which binds to the nuclear receptors RAR and RXR to control transcriptional expression of numerous target genes. Among its recently recognized effects on immunity, retinol modulates the maturation, antigen presentation, migration, and T-cell priming functions of dendritic cells [41–43], as well as the generation of T-regulatory cells and suppression of Th17 cells [44]. Retinoic acid has also been shown to be crucial to the maintenance of the Th1 lineage [45], an essential component of an effective immune response to tuberculosis.

In addition to its impact on acquired immunity, recent evidence suggests that vitamin A may play a role in innate immune responses to Mtb. Wheelwright et al. found that all-trans-retinoic acid exerts an antimicrobial effect on Mtb in human monocytes through its effect on cellular cholesterol efflux mediated through the NPC2 gene [6]. Stimulation of primary human monocytes with all trans-retinoic acid increased expression of NPC2 and reduced cellular cholesterol through a mechanism that is distinct from the antimicrobial activity of 1,25-dihydroxyvitamin D3 [6]. Mtb has been shown to use host lipids as a nutrient source, and host cholesterol mediates Mtb persistence and survival within macrophages [46]. Taken together, these findings suggest that low vitamin A levels may impair host’s ability to control TB infection after exposure.

Our study had notable limitations. Participants were followed for one year, a relatively short period of time given the slow pathogenesis of tuberculosis. Because we had relatively few blood samples from children under ten, we were unable to assess the impact of vitamin A deficiency in that age group. Almost all our study participants had been vaccinated with BCG so we are unable to distinguish between the possibilities that vitamin A enhanced BCG-induced adaptive immunity or provided direct protection against TB progression. Although we found no evidence of an interaction between vitamin A and vitamin D on TB disease risk, our study may not have been powered to detect such an interaction.

Most importantly, despite our efforts to ensure that nutritional assessments occurred prior to the development of TB disease, we could not rule out the possibility that participants had unrecognized TB disease at baseline, which may have led to the low retinol levels we observed. We addressed this issue by conducting two sensitivity analyses, one of which stratified on baseline infection status and one which excluded cases diagnosed within 90 days of index patient enrollment. In both analyses, risk of TB disease remained inversely associated with vitamin A quartiles regardless of baseline TB infection status or timing of diagnosis. Nevertheless, since active TB may remain clinically undetected for months, especially in children, and since acute infections are known to affect serum vitamin A levels, there is no way to rule out reverse causality without conducting a clinical trial of vitamin A supplementation in people at high risk for TB.

In conclusion, we find that vitamin A levels among persons exposed at home to a TB patient strongly predicted incident TB disease within 12 months of follow-up in a dose-dependent manner. Our data raise the possibility that screening for vitamin A levels among people at high risk for TB could help identify those likely to develop disease in the near future so they could be targeted for early intervention. In the event that the association between VAD and tuberculosis progression proves to be causal, routine vitamin A supplementation among persons at high risk for developing TB disease (e.g. household contacts of TB cases or individuals in congregate settings such as prisons) may provide an inexpensive, safe and effective means of preventing progression from TB infection to TB disease.

